# Special care is needed in applying phylogenetic comparative methods to gene trees with speciation and duplication nodes

**DOI:** 10.1101/719336

**Authors:** Tina Begum, Marc Robinson-Rechavi

## Abstract

How gene function evolves is a central question of evolutionary biology. It can be investigated by comparing functional genomics results between species and between genes. Most comparative studies of functional genomics have used pairwise comparisons. Yet it has been shown that this can provide biased results, since genes, like species, are phylogenetically related. Phylogenetic comparative methods should allow to correct for this, but they depend on strong assumptions, including unbiased tree estimates relative to the hypothesis being tested. Such methods have recently been used to test the “ortholog conjecture”, the hypothesis that functional evolution is faster in paralogs than in orthologs. Whereas pairwise comparisons of tissue specificity (*τ*) provided support for the ortholog conjecture, phylogenetic independent contrasts did not. Our reanalysis on the same gene trees identified problems with the time calibration of duplication nodes. We find that the gene trees used suffer from important biases, due to the inclusion of trees with no duplication nodes, to the relative age of speciations and duplications, to systematic differences in branch lengths, and to non-Brownian motion of tissue-specificity on many trees. We find that incorrect implementation of phylogenetic method in empirical gene trees with duplications can be problematic. Controlling for biases allows to successfully use phylogenetic methods to study the evolution of gene function, and provides some support for the ortholog conjecture using three different phylogenetic approaches.

## Introduction

The “ortholog conjecture”, a standard model of phylogenomics, has become a topic of debate in recent years (Koonin 2005; Studer and Robinson-Rechavi 2009; Nehrt et al. 2011; Altenhoff et al. 2012; Chen and Zhang 2012; Gabaldón and Koonin 2013; Rogozin et al. 2014; Kryuchkova-Mostacci and Robinson-Rechavi 2016; Dunn et al. 2018; Stamboulian et al. 2020). The ortholog conjecture is routinely used by both experimental and computational biologists in predicting or understanding gene function. According to this model, orthologs (i.e. homologous genes which diverged by a speciation event) retain equivalent or very similar functions, whereas paralogs (i.e. homologous genes which diverged by a duplication event) share less similar functions (Studer and Robinson-Rechavi 2009). This is linked to the hypothesis that paralogs evolve more rapidly. This hypothesis was challenged by results suggesting that paralogs would be functionally more similar than orthologs (Nehrt et al. 2011). Such findings not only raised questions on the evolutionary role of gene duplication but also questioned the reliability of using orthologs to annotate unknown gene functions in different species (Sonnhammer et al. 2014). Several studies (Altenhoff et al. 2012; Chen and Zhang 2012; Rogozin et al. 2014; Kryuchkova-Mostacci and Robinson-Rechavi 2016) later found support for the ortholog conjecture, mostly based on comparisons of gene expression data.

While all previous studies of the ortholog conjecture had used pairwise comparisons of orthologs and paralogs, a recent article suggested that this was flawed, and that phylogenetic comparative methods should be used (Dunn et al. 2018). Phylogenetic structure can violate the fundamental assumption of independent observations in statistics, and thus ignoring it can lead to mistakes (Felsenstein 1985). A solution is to use phylogeny-based methods. Phylogenetic Independent Contrast (PIC) (Felsenstein 1985), and Phylogenetic Generalized Least-Square (PGLS) (Martins and Hansen 1997; Grafen 1989; Rohlf 2001) are the most commonly used phylogenetic comparative methods. They were developed under a purely neutral model of evolution, i.e. Brownian motion (BM). Such Brownian process have been extended using a maximum likelihood approach, to allow for different rates of evolution on different branches of a phylogeny (O’Meara et al. 2006; Thomas et al. 2006), and to include stabilizing selection in which the trait is shifted towards a single fitness optimum, or multiple different adaptive optima (i.e. “Ornstein-Uhlenbeck” or OU process) (Hansen 1997; Butler and King 2004; Beaulieu et al. 2012). These phylogenetic data modeling with different modes of trait evolution (e.g. BM, OU) require a priori knowledge of different states on the tree. Other approaches implemented a Markov chain Monte Carlo (MCMC) sampling in a Bayesian framework to accurately estimate the number, location, and magnitude of shifts in evolutionary rates, or in optimal trait values without a priori assignment of states (Eastman et al. 2011; Pennell et al. 2014; Uyeda and Harmon 2014; Catalan et al. 2019). Bayesian approaches are time consuming, while OU modeling with phylogenetic lasso algorithm allows a faster detection of directional selection due to a shift in optimal trait value (Khabbazian et al. 2016). Moreover, OU has been used to model gene expression evolution (Rohlfs and Nielsen 2015; Chen et al. 2019). Among all the phylogenetic methods, PIC is widely adopted for its relative simplicity, and its applicability to a wide range of statistical procedures (Cooper et al. 2016a; Dunn et al. 2018). The performance of PIC relies on three basic assumptions: a correct tree topology; accurate branch lengths; and trait evolution following Brownian motion (where trait variance accrues as a linear function of time) (Felsenstein 1985; Garland 1992; Garland et al. 1992; Díaz-Uriarte and Garland 1998; Freckleton and Harvey 2006; Cooper et al. 2016a). If any of these assumptions is incorrect, this can lead to incorrect interpretation of results without control for biases (Diaz-Uriarte and Garland 1996; Díaz-Uriarte and Garland 1998). While previous applications of PIC used multivariate traits on pure speciation trees to explore the relationship between them, Dunn et al. (2018) took an innovative approach in applying PIC to compare the divergence rates of a univariate trait between two different node events (“speciation” and “duplication”), to test the ortholog conjecture. They performed extensive analyses in support of their results. However, such an application might be problematic since the time of occurrence of gene duplication, one of the two types of events compared, is unknowable by external information (e.g. no fossil evidence). Therefore, further study is required to understand why Dunn et al. (2018) obtained results which are inconsistent with previous studies. It is possible that all the conclusions drawn by previous studies on gene duplication are incorrect due to overlooking phylogenetic tree structure. If so, it should be well supported.

We re-examined the data of Dunn et al., after reproducing their results using the resources and scripts provided by the authors (Dunn et al. 2018). We have uncovered problems with the use of PIC on biased calibrated gene trees, violation of the underlying assumptions, and the inclusion of pure speciation gene trees. We used PIC on gene trees after fixing calibration bias for old duplication nodes. With proper controls, the phylogenetic method supports the ortholog conjecture. To verify this result, we also applied data modeling approaches using a maximum likelihood framework, and using a reversible-jump Bayesian MCMC algorithm. Support for the ortholog conjecture still holds with proper controls.

## Results

### Issues with straightforward application of Phylogenetic Independent Contrasts (PICs)

Dunn et al. (2018) have made a relevant argument that the test should be done in a phylogenetic framework, since closely related species or genes tend to share more similar traits. They applied PIC method to a processed dataset of 8520 time-calibrated trees (details in the Materials and Methods, Table 1) by assuming that the computed node contrasts (PICs) are always phylogenetically independent, and reported evidence in contradiction with the ortholog conjecture for tissue-specificity *τ* (median: PIC_speciation_ = 0.0072, PIC_duplication_ = 0.0051, one-sided Wilcoxon test *P* = 1). Yet the same data supported the ortholog conjecture when analyzed by pairwise comparisons, both in Kryuchkova-Mostacci and Robinson-Rechavi (2016), and in the re-analysis by Dunn et al. (2018). To understand the incongruence between results of PIC and of pairwise comparison approaches, they performed simulations of *τ* on their trees under the OC (ortholog conjecture), and under a null of uniform Brownian motion. PICs and pairwise comparisons have different expectations under the null (*σ*^2^_duplication_ = *σ*^2^_speciation_) and under the ortholog conjecture (*σ*^2^_duplication_ > *σ*^2^_speciation_) (Supplementary fig. S1). The simulation results of Dunn et al. (2018) indicated that the pairwise comparisons of events could not distinguish the two scenarios (null and OC), unlike the PIC method. As the result on their empirical data resembled their null simulation result, they questioned both the use of pairwise comparisons, and the support for the ortholog conjecture from tissue specificity data.

**Table 1:** Information on different tree sets, number of internal node events, and node ages used in this reanalysis.

| Datasets | Number of trees | Number of speciation events | Number of duplication events | Number of NA events | Maximum speciation node age (My) | Maximum duplication node age (My) |
| --- | --- | --- | --- | --- | --- | --- |
| Dunn et al.: full set | 8520 | 67911 | 21071 | 26794 | 296 | 11799977 |
| Dunn et al.: trees with strong phylogenetic signals | 2082 | 13118 | 4056 | 5186 | 296 | 1342 |
| This study: after excluding pure speciation trees | 4288 | 38882 | 15274<br>(8556 young + 6718 old) | 15201 | 296 | 1175 |

*Note-* My: Million years; young: age ≤ 296 My; old: age $>$ 296 My.

To understand their results, we first reproduced and reanalyzed the data of Dunn et al. (2018) by focusing on the phylogenetic approach. Dunn et al. reported a non-significant result (*P* = 1) for the PIC under the null simulation as well as for the empirical data, using a Wilcoxon one-tailed rank test to check if the contrasts of duplication events are higher than the contrasts of speciation events. Surprisingly, our reanalysis with a Wilcoxon two-tailed rank test on the same data shows that the PIC rejects the null hypothesis on the null simulations (Fig. 1A), with significant support for higher contrasts after speciation than duplication. This means that the PIC method supports a trend opposite to the trend expected under the ortholog conjecture in a null simulation. This was robust to repeating the simulations with different random seed number (Supplementary fig. S2). This indicates that neither of the approaches, PIC or pairwise, worked properly for these calibrated trees, since both the approaches reject the null hypothesis when simulations are performed under the null. Similarly, when we used a Wilcoxon two-tailed rank test instead of a one-tailed test on the empirical data, the non-significant result (*P* = 1) (Dunn et al. 2018) was also significant (*P* < 2.2e^-16^) in the same unexpected direction as the null simulation results.

**Figure 1:**
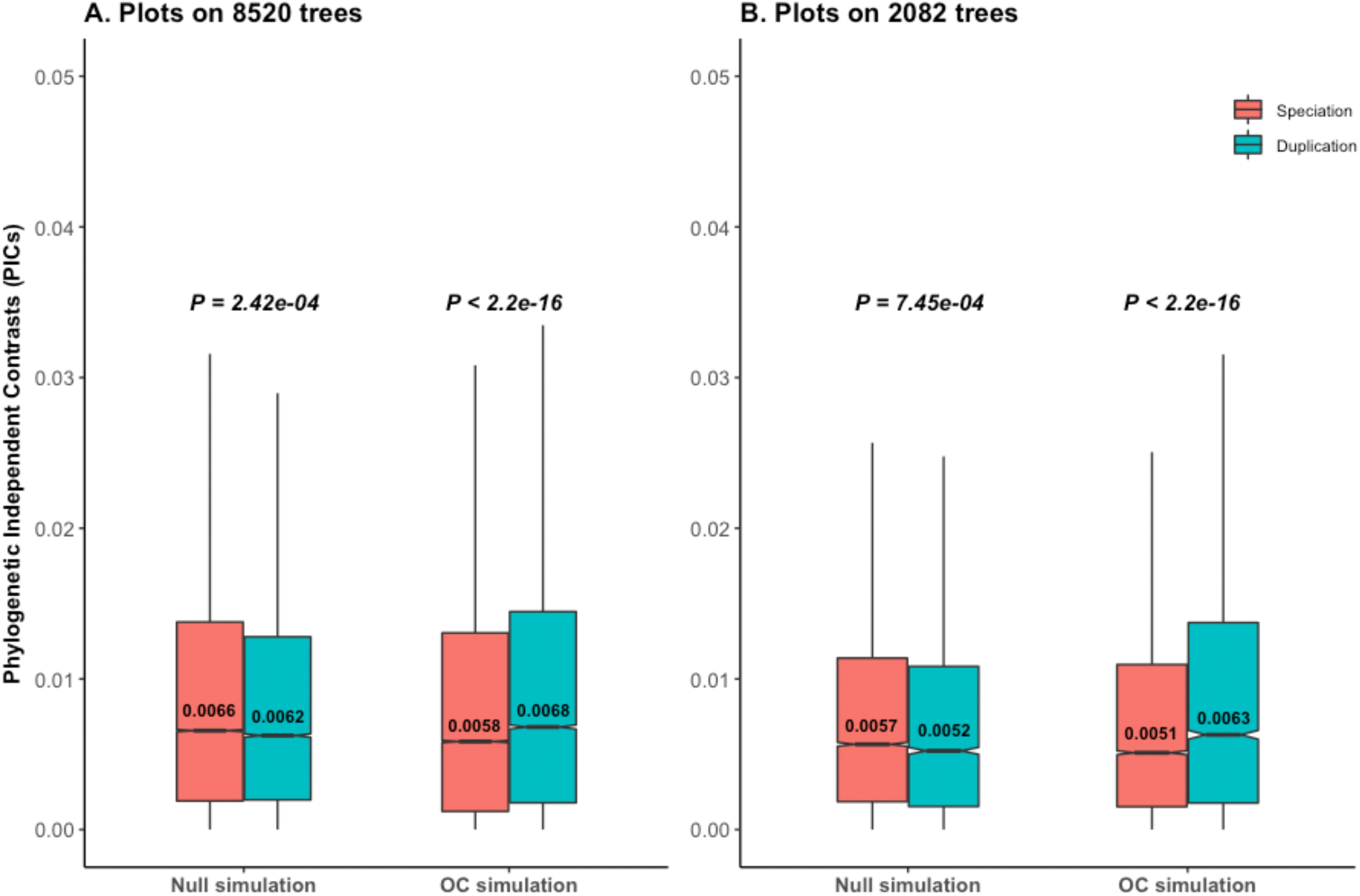
Reanalyses of phylogenetic simulation data of Dunn et al. (2018). *P* values are from Wilcoxon two-tailed tests. Values inside boxplots denote median PIC values of the corresponding events. In null simulations, there should be no difference in contrasts between events. In OC (Ortholog Conjecture) simulations, contrasts are expected to be higher for duplication than for speciation. (A) Higher contrasts for speciation than duplication reject the null hypothesis under null simulation scenario for all empirical time calibrated gene trees. (B) Results are similar with a subset of trees with strong phylogenetic signal for *τ*.

Statistical non-independence among species trait values because of their phylogenetic relatedness can be measured by phylogenetic signal (Pagel 1999; Freckleton et al. 2002; Blomberg et al. 2003; Münkemüller et al. 2012; Molina-Venegas and Rodríguez 2017). Use of the PIC is mainly important for the data sets with strong phylogenetic signal, where it allows to recover phylogenetically independence. Dunn et al. (2018) used Blomberg’s K. Its value ranges from 0 to ∞ for each tree, where a value of 0 indicates no phylogenetic signal for the trait studied, and a value close to 1 or higher indicates strong phylogenetic signal (Pagel 1999; Freckleton et al. 2002; Blomberg et al. 2003; Münkemüller et al. 2012; Molina-Venegas and Rodríguez 2017). With a cutoff of K > 0.551, Dunn et al. (2018) obtained only 2082 trees (Table 1), 24.4% of the total, with strong phylogenetic signal. The phylogenetic method still rejects the null hypothesis under null simulations for those 2082 trees using a Wilcoxon two-tailed rank test (Fig. 1B), showing that the problem is not simply due to low phylogenetic signal. Using a cut-off of *P* < 0.05 together with K > 0.551 leads to 1135 statistically significant trees with strong phylogenetic signals, for which we obtained a similar result (Supplementary fig. S3). This means that the bias is not limited to the selection of tree sets or to the number of speciation or duplication events used for the analyses. Since the trend was similar for these 1135 trees we continued analyses with the 2082 trees of Dunn et al. (2018) for consistency.

The accuracy and performance of the PIC method largely depend on proper branch length calibration in absolute time (e.g. in Million Years – My) (Garland 1992; Díaz-Uriarte and Garland 1998; Cooper et al. 2016a). We thus investigated possible biases created during calibration of gene trees. Due to non-availability of external references for duplication time points (e.g. no fossils), Dunn et al. (2018) used only 7 speciation time points to calibrate gene trees. The ages of other node events are estimated using the penalized likelihood method (Sanderson 2002) by the chronos() function of the “ape” R package (Paradis et al. 2004), and varies for the same duplication clade labels even within the same gene trees. The oldest speciation age for their calibrated trees was 296 My (Table 1), corresponding to the use of chicken as the outgroup. Surprisingly, the calibrated node age of the oldest duplication event was 11799977 My (Table 1, Supplementary table S1), that is, 2600 times older than the Earth. This is indicative of issues with calibration. The tree pruning to species with *τ* data (details in Materials and Methods) lead to trees for which all nodes older than 296 My are duplication or NA events, even if there were older speciation events present before pruning (Supplementary fig. S4A). If the root node of a pruned tree is a speciation, the duplication ages are constrained by speciation ages. Otherwise, there are no constraints for the duplication events older than the oldest speciation events (Supplementary fig. S4, Supplementary Table S1), which can introduce a calibration bias. This unreliable branch length estimation for the old duplication nodes eventually led to much larger expected variances for gene duplication events than for speciation events (Supplementary figs. S5A and S5B).

PIC of a node is a ratio of changes in trait values (*τ* here) for descendant nodes to their expected variance, i.e. the lengths of the two branches that connect the node to its two descendants. This means that similar changes in *τ* for two nodes can produce different PIC values, with the lower contrast for the node with higher expected variance (i.e., calibrated branch length). In the null simulations only the *τ* values are simulated, while the branch lengths (hence the expected variances) are taken from the empirical data, and thus share its biases. This explains why contrasts are lower for duplications than for speciations under null simulations as well as with empirical data. Such calibration bias in branch lengths violates the second assumption of PIC applicability, and inflates type I error rates (Diaz-Uriarte and Garland 1996; Díaz-Uriarte and Garland 1998).

### Randomization tests to assess the performance of phylogenetic method

We used randomization tests to assess bias in different analyses of the empirical dataset. Our expectation is that the trend of the empirical result should differ from the randomized ones. In a first randomization test, we permuted the *τ* values across the tips of each tree without altering the node events of the trees. By such randomization, the real phylogenetic relationships between trait values are removed for each tree. When we compared the node contrasts of the speciation and duplication events computed based on these 8420 randomized *τ* trees (Fig. 2A), we found the same pattern as reported for the empirical gene trees by Dunn et al. (2018), contrary to expectation. It confirms that results are driven by their large differences in branch lengths (i.e. in expected variances) (Fig. 2B), as on simulated null data. Any effect of trait divergence rates of speciation and duplication events is always masked by this branch length difference of node events. This violates the basic assumption of applicability of the PIC method to Brownian trait evolution. To remove the problem of difference in expected variances of the two events, we performed a second randomization test: we kept the original *τ* value for tips but randomly shuffled the events (duplication, speciation, or NA) of internal nodes of the 8420 empirical gene trees to maintain the original proportions of speciation and duplication events. The resulting trend (Fig. 2C) still resembled the empirical gene trees data. This appears due to the fact that the majority of the nodes are speciations (Fig. 2D, Table 1) with node ages ≤ 296 My. Most of the trees with many duplication events on the other hand have ancient duplication events for which the evolutionary rates of duplication are often masked by the effect of longer branch lengths. Opposite to our expectation, the calibrated trees with no or few duplications have higher overall nodes contrast (apparent fast evolution) than trees with many duplications (apparent slow evolution). This might be due to greater difficulty in detecting paralogs for fast evolving genes. Therefore, reshuffling of the events may not change the observed pattern of higher speciation contrasts than duplication contrasts.

**Figure 2:**
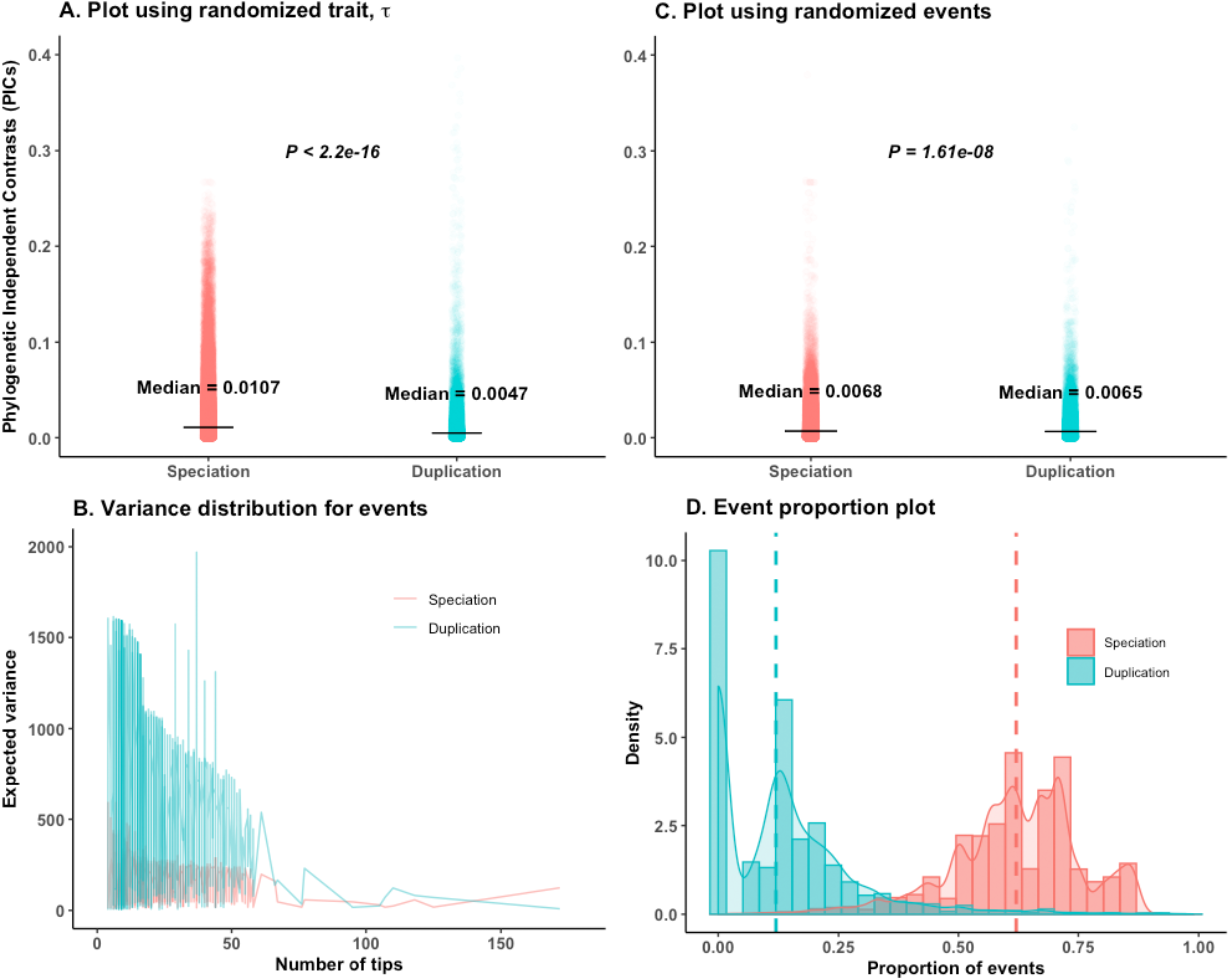
Analyses on calibrated empirical gene trees of Dunn et al. (2018). *P* values are from Wilcoxon two-tailed tests. (A) Randomly shuffling the *τ* values of the tips for 8520 gene trees does not alter the empirical trend of an opposite trend to the ortholog conjecture. (B) The expected variance is much higher for duplication than speciation events irrespective of the number of tips considered for the study. (C) Using the original *τ* data, if we permute the events (Speciation or Duplication or NA) of the nodes, the trend of result remains. (D) The proportions of speciation events is much higher than duplication events for all time-calibrated trees; the dotted line represents the median proportion of both events; a high proportion of trees have no duplication events.

Out of 8520 calibrated trees, 2990 were pure speciation trees with no duplication events. For these 2990 trees, random shuffling of events had no impact. To avoid this bias, we removed those 2990 speciation trees as well as trees with negative branch lengths, and randomized the trait or the internal node events 100 times on the remaining 5479 trees. However, we still always obtained significantly higher contrasts of speciation than of duplication (Supplementary figs. S6A and S6B). The randomization tests pattern is the same when we used 2082 trees with strong phylogenetic signals (Supplementary figs. S6C and S6D).

All these analyses indicate that the results reported by Dunn et al. (2018) are biased by the calibrated phylogeny structures, and that this bias is not easy to correct. We propose three approaches to correct for this bias and recover a proper phylogenetic signal of trait evolution.

### Approach-1: PIC with diagnostic tests

Diagnostic tests (details in the Materials and Methods) for each tree are essential to ensure phylogenetic independence of node contrasts, especially since there is evidence of bias in the calibrated trees. This can be verified by the lack of correlation between the absolute value of PICs of *τ* and their standard deviations, node height, node age, or node depth (Garland 1992; Garland et al. 1992; Diaz-Uriarte and Garland 1996; Díaz-Uriarte and Garland 1998; Freckleton 2000; Freckleton and Harvey 2006; Cooper et al. 2016a). A statistically significant negative or positive correlation in any of the diagnostic tests confirms that the PICs for that tree are non-independent (Garland 1992; Garland et al. 1992; Diaz-Uriarte and Garland 1996; Díaz-Uriarte and Garland 1998; Freckleton 2000; Freckleton and Harvey 2006; Cooper et al. 2016a); in practice, we used *P* < 0.05 for significance.

We performed such diagnostic tests on 4288 trees, for which calibration biases are fixed for old duplication nodes (see Materials and Methods, Table 1). Among them only 2088 (48.7%), which includes 15321 speciation and 6213 duplication nodes, passed all 4 diagnostics tests for *τ* evolution. We performed our PIC analyses separately for 3948 young (≤ 296 My, the oldest speciation in the trees) and 2265 old (> 296 My) duplication events. Analyses on young duplicates after diagnostic tests provided support for the ortholog conjecture (Fig. 3), but old duplicates did not. Randomization tests showed patterns distinct from real data only for the young duplicates (Supplementary figs. S7A and S7B), indicating a biological pattern rather than a data bias. Thus PIC on the trees after diagnostic plot tests supports the ortholog conjecture for young duplicates, whereas the inference remains biased for older duplicates.

**Figure 3:**
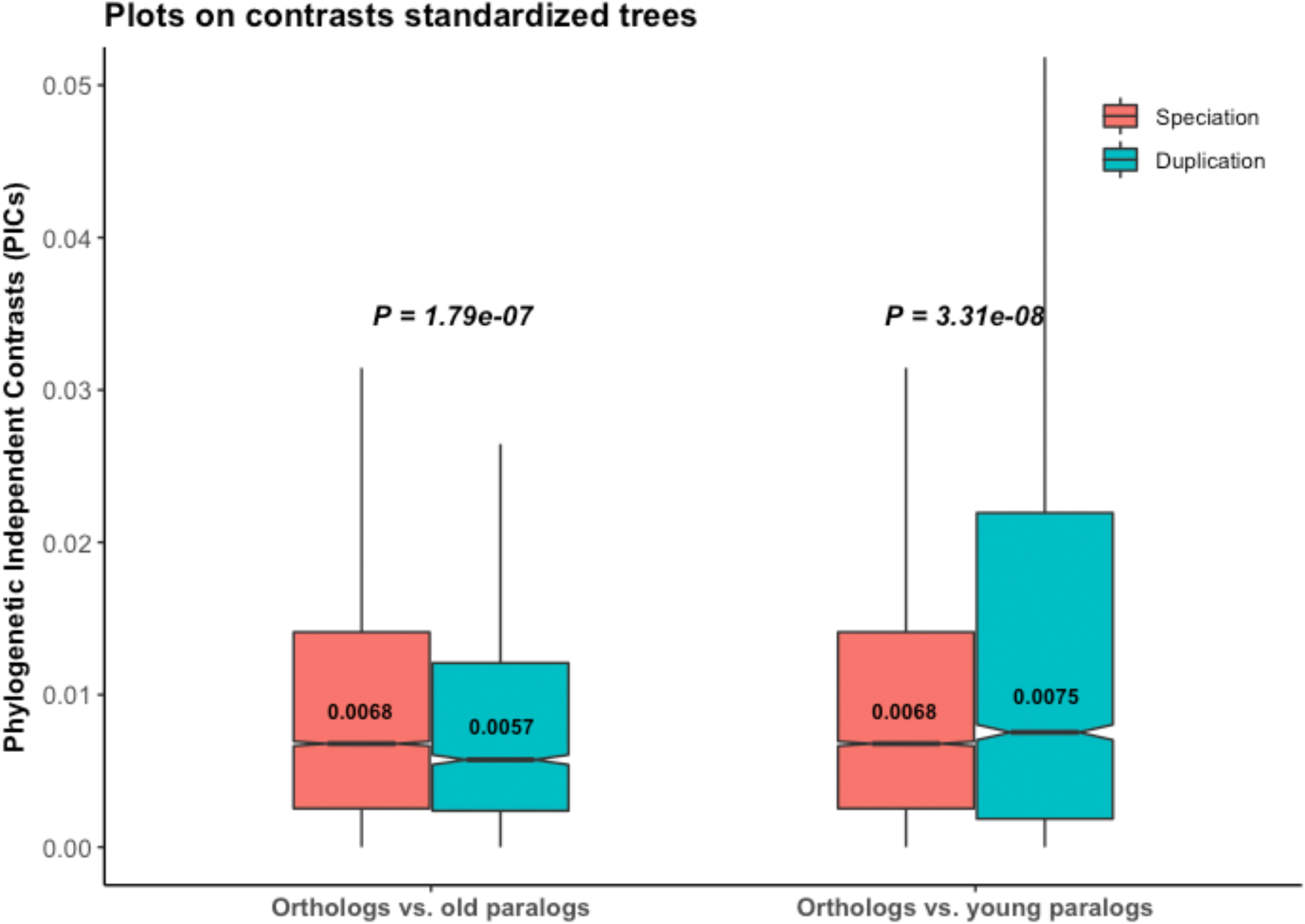
The ortholog conjecture test on *τ* for trees passing diagnostic plot tests. *P* values are from Wilcoxon two-tailed tests. Values inside boxplots denote median PIC values of the corresponding events. Young duplicates: age ≤ 296 My, the maximum speciation age; old duplicates: age > 296 My.

### Approach-2: PIC with branch length transformation

Most phylogenetic methods are developed for the Brownian model of trait evolution, including the PIC method (Felsenstein 1985; Cornwell and Nakagawa 2017). Deviations from pure BM violate the fundamental assumptions of PIC applicability and can affect its performance for testing hypotheses about correlated evolution (Garland 1992; Garland et al. 1992; Diaz-Uriarte and Garland 1996; Díaz-Uriarte and Garland 1998). Using model-fitting (see Materials and Methods), we found that 75.6% gene trees (Supplementary fig. S8) supported the Ornstein-Uhlenbeck (OU) model. Remedial measures such as branch length transformations along with diagnostic tests, can substantially recover the performance of the PIC methods when character evolution is not BM or when contrasts are non-independent of the phylogeny (Garland et al. 1992; Diaz-Uriarte and Garland 1996; Díaz-Uriarte and Garland 1998).

We applied branch length transformation (details in the Materials and Methods) on all 4288 trees, along with diagnostic tests for consistency. We found substantial support for the ortholog conjecture for the 4190 trees (97.7%) which pass diagnostic tests after branch length transformation (Fig. 4A). Due to the lack of absolute age for these transformed trees, we did not distinguish young and old duplicates. Applying such branch length transformation then diagnostic tests to the gene trees of Dunn et al. we also found support for the ortholog conjecture in 98.8% (8417 out of 8520) (Supplementary fig. S9A), as well as for 99.9% (2080 out of 2082) of their trees with strong phylogenetic signal (Supplementary fig. S10A). Randomization tests on all these sets of trees following branch length transformations clearly showed distinct patterns compared to the empirical data (Figs. 4B and 4C, Supplementary figs. S9B, S9C, S10B, S10C), indicating that results are not due to inference bias once the data is properly transformed.

**Figure 4:**
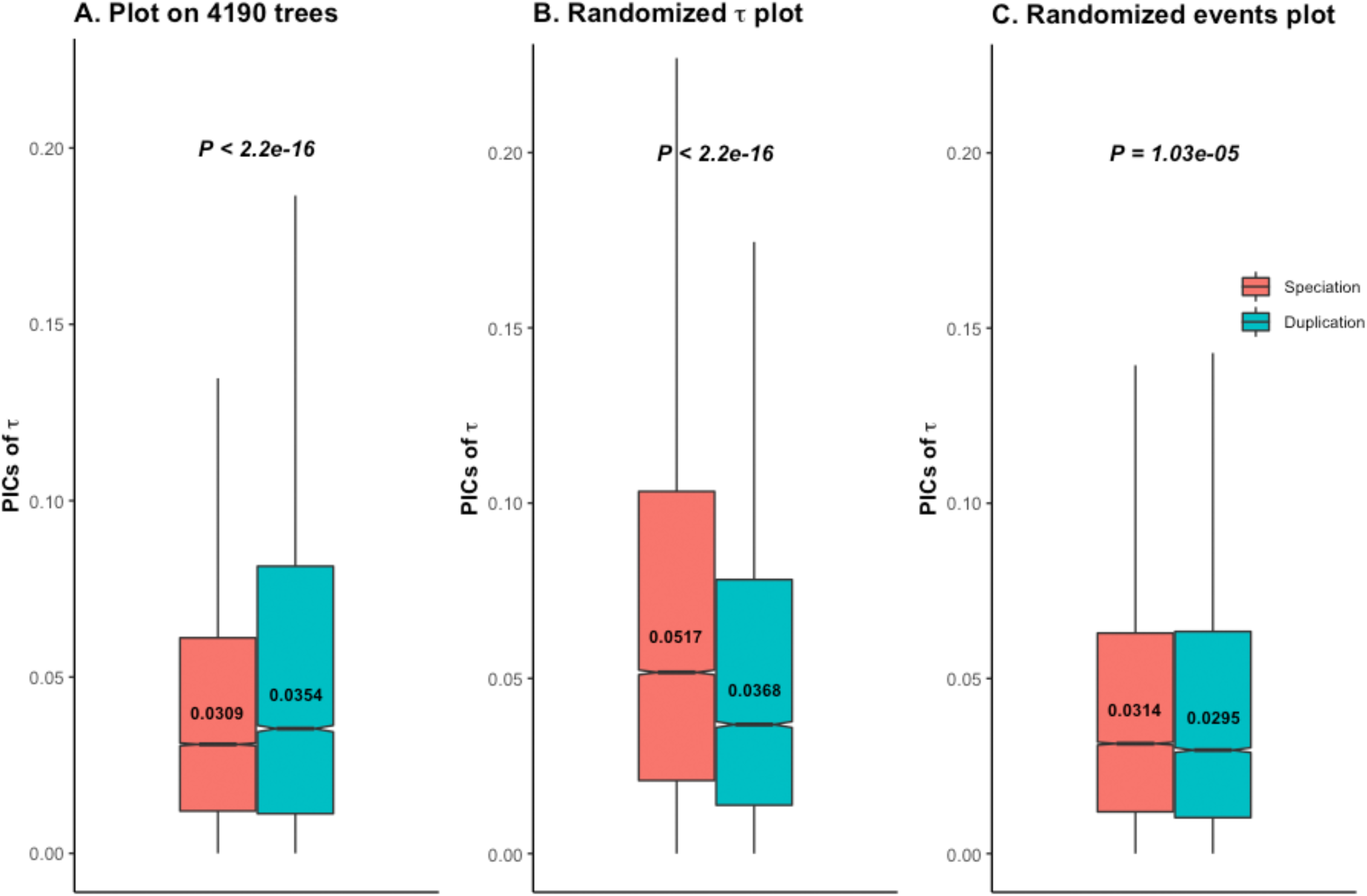
The ortholog conjecture test for contrasts standardized branch transformed trees. *P* values are from Wilcoxon two-tailed tests. Values inside boxplots denote median PIC value of the corresponding event. (A) Using 4190 out of 4288 calibrated trees that passed diagnostic tests following branch length transformation. (B) Permuting *τ*, and (C) permuting internal events on contrasts standardized branch length transformed trees produces distinct patterns compared to the empirical gene trees of (A).

### Approach-3: Phylogenetic data modeling

State dependent model-fitting allows to compare the evolutionary rates (*σ*^2^), and the changes in adaptive optimum value (θ) associated to specific states (speciation or duplication) for each tree (Beaulieu et al. 2012; Clavel et al. 2015). Under the ortholog conjecture, our expectation is that there should be more shifts in optimum value of *τ* between paralogs than orthologs. Moreover, the evolutionary rates after duplication should be higher than after speciation (*σ*^2^_duplication_ > *σ*^2^_speciation_). Of course, trends on empirical data should differ from randomized ones. When we modeled the evolution of *τ* (see Materials and Methods), 32 out of 4288 trees failed to fit any model due to invariance in τ. Among the others, 308 supported BM1, 704 BMM, 2874 OU1, and 370 OUM, as the best fit models (Supplementary fig. S8). We performed our analyses separately for young and old duplicates.

On the 8.6% multi optima trees (OUM) the optimum value are significantly higher for both young and old duplications (θ_dup_ > θ_spe_) (Supplementary Table S2). Thus paralogs regime shift towards higher tissue-specificity. These results are not observed on randomized trees, supporting a biological pattern in the data (Supplementary Table S2).

We also applied a Bayesian method (Udeya and Harmon 2014) on them to quantify the number of adaptive optimum shifts, as suggested for small trees (Cooper et al. 2016b). Unlike the other approach, such detection of evolutionary shifts in a phylogeny does not need a priori knowledge of different states on the tree. Using a strict posterior probability threshold of ≥ 0.7 with this method, we find that most optimum shifts per branch for *τ* follow duplications (median after speciation: 0%, after duplication: 12.5%, paired two-sided Wilcoxon rank-sum test *P* < 2.2e^-16^). An OU model can often be incorrectly favored over a BM model in a maximum likelihood framework when applied to trees with < 200 tips (Cooper et al. 2016b). Our gene trees have a median of only 15 tips. We thus applied a conservative Bayesian approach on all of the 3244 trees for which OU was the preferred model (OU1 + OUM). Even with such a strict posterior probability threshold of ≥ 0.7, 1101 trees (33.9%) still supported the OUM model, including 901 trees identified as OU1 by maximum likelihood. We detected the same trend of optimum shifts per branch (median after speciation: 2.3%, after duplication: 10%, paired Wilcoxon rank-sum test *P* < 2.2e^-16^). These results are largely consistent for both young and old duplicates (Table 2; Supplementary Table S3). However, the rates of optimum shifts are faster only for young duplicates (Table 2; Supplementary Table S3).

**Table 2:** Summary statistics on 1101 OUM trees passing a posterior probability cutoff of ≥ 0.7 in a Bayesian framework.

| Duplication<br>Age | Proportions of regime shifts per branch |  | Paired two-sided<br>Wilcoxon rank sum test | Regime shift rates (shifts/My) |  | Two-sided<br>Wilcoxon rank test |
| --- | --- | --- | --- | --- | --- | --- |
|  | After speciation | After duplication |  | After speciation | After duplication |  |
| Young | 3.1% | 4.5% | $3.4e^{-12}$ | 0.013 | 0.031 | $1.7e^{-11}$ |
| Old | 2.6% | 10% | $< 2.2e^{-16}$ | 0.013 | 0.0023 | $< 2.2e^{-16}$ |

*Note-* Above analyses include 13824 speciation, 3027 young and 2814 old duplication events. Values shown in the table indicate median values. The difference in proportions of regime shifts per branch after speciation events for two types of duplications is due to the different sets of trees used. Few trees shared both types of duplicates. Proportions of regime shifts per branch of events is estimated for each tree, and thus paired Wilcoxon test is used to compare the difference. A single gene tree can have multiple optima shift rates for events, and thus two-sided Wilcoxon rank test was used for comparison.

Analyses on the trees where σ^2^ varies between events (BMM) also supports the ortholog conjecture for young duplicates (Table 3). Randomized data showed distinct patterns from empirical data. However, again there was neither support for the ortholog conjecture nor signal relative to randomization for the old duplicates.

**Table 3:** Summary statistics for Brownian trees.

| Duplication age | Data | $\sigma^2_{\text{Speciation}}$ | $\sigma^2_{\text{Duplication}}$ | $\sigma^2_{\text{Duplication}} / \sigma^2_{\text{Speciation}}$ | *P*-value |
| --- | --- | --- | --- | --- | --- |
| Young | Empirical<br>(n<sub>Speciation</sub>= 4642;<br>n<sub>Duplication</sub> = 1742) | 9e<sup>-5</sup> | 1.4e<sup>-4</sup> | 1.5 | 5e<sup>-12</sup> |
| | Randomized $\tau$<br>(n<sub>Speciation</sub>= 4618;<br>n<sub>Duplication</sub> = 1723) | 6.9e<sup>-4</sup> | 2.2e<sup>-4</sup> | 0.32 | 1.4e<sup>-13</sup> |
|  | Randomized events<br>(n<sub>Speciation</sub>= 3215;<br>n<sub>Duplication</sub> = 1438) | 1.7e<sup>-4</sup> | 8.5e<sup>-5</sup> | 0.5 | 0.02 |
| Old | Empirical<br>(n<sub>Speciation</sub>= 5356;<br>n<sub>Duplication</sub> = 1295) | 1.7e<sup>-4</sup> | 2e<sup>-9</sup> | 1.2e<sup>-5</sup> | < 2.2e<sup>-16</sup> |
| | Randomized $\tau$<br>(n<sub>Speciation</sub>= 5337;<br>n<sub>Duplication</sub> = 1291) | 9.1e<sup>-4</sup> | 2.5e<sup>-10</sup> | 2.7e<sup>-7</sup> | < 2.2e<sup>-16</sup> |
|  | Randomized events<br>(n<sub>Speciation</sub>= 2788;<br>n<sub>Duplication</sub> = 800) | 1.8e<sup>-4</sup> | 2.1e<sup>-9</sup> | 1.2e<sup>-5</sup> | < 2.2e<sup>-16</sup> |

*Note:* Median values of $\sigma^2$ are shown. *P*-value from paired two-sided Wilcoxon test.

## Discussion

We agree with Dunn et al. (2018) that evolutionary comparisons should be done considering a phylogenetic framework when possible. However, this does not imply that phylogenetic methods can be applied easily to phylogenomics. To get a clear picture, we limited our study to the same gene trees used by Dunn et al. (2018). Our reanalysis identified problems generated by the time calibration of old duplication nodes of pruned trees, the inclusion of pure speciation gene trees, and violations of the Brownian model. The strongest bias was for duplication nodes preceding the oldest speciation nodes. This, in turn, introduced several biases in the analyses, and influenced results.

When we identified and controlled for such biases, PIC results changed to support the ortholog conjecture, consistent with our previous pairwise analysis (Kryuchkova-Mostacci and Robinson-Rechavi 2016) on the same *τ* data. Our fundamental point is that the conclusions drawn by Dunn et al, but also by anyone else who will have followed the same approach of applying PIC to gene trees, are not reliable unless extreme care is taken. This is because gene trees with orthologs and paralogs have more complex evolutionary histories, and different sampling biases, than species trees for which these methods were developed.

To date, a few studies have applied phylogenetic comparative methods to understand the effect of gene duplication on functional evolution (Oakley et al. 2005; Oakley et al. 2006; Eng et al. 2009; Rohlfs and Nielsen 2015; Dunn et al. 2018; Fukushima and Pollock 2020). None before Dunn et al. applied PIC method to compare speciation and duplication events on the same trees using a single continuous trait. Such application requires thorough testing of the fundamental assumptions of the method on such time calibrated trees (Garland 1992; Garland et al. 1992; Diaz-Uriarte and Garland 1996; Díaz-Uriarte and Garland 1998; Freckleton 2000; Freckleton and Harvey 2006; Cooper et al. 2016a). Hence, we explored whether the application of a phylogenetic method might inflate errors (e.g. rejection of the null hypothesis in null condition) if applied without assumption testing. Indeed, it is the case (Figs. 1A and 1B). Along with the calibration bias for old duplication nodes, the relative ages of the speciation and duplication events strongly differ in these trees due to the choice of species. Using such trees without control for biases may bring about lack of statistical power to detect the signal of ortholog conjecture, and even bias towards an opposite pseudo-signal.

Time calibration of ancient duplication events is one of the major issues we uncovered. The approach of Dunn et al. considered pruned trees with available trait (*τ* here) data for time-calibration using speciation time points (see Materials and Methods). Such pruned trees often have many duplication nodes older than the oldest speciation nodes. Sequence based evolutionary rate (e.g., dN/dS) analyses in different species have found higher sequence evolutionary rate following gene duplication (Conant and Wagner 2003; Kim and Yi 2006; Scannell and Wolfe 2008; Han et al. 2009; Studer and Robinson-Rechavi 2009; Panchin et al. 2010; Pegueroles et al. 2013; Pich and Kondrashov 2014; Holland et al. 2017). Therefore, calibration bias is not surprising for those duplication nodes in the absence of time constraints (Supplementary figs. S4A-S4C, Supplementary Table S1). Instead, we performed time calibration before pruning, so that the oldest speciation time points can provide upper age limits and reduce calibration bias (Supplementary figs. S4D-S4F). This is strongly recommended since the performance of the phylogenetic methods rely on accurate branch length information, especially for multi-states univariate trait analysis.

Dunn et al. (2018) performed several analyses (e.g. added random noise in the speciation calibration time points, extended terminal branch length, removed old duplication nodes, etc.) to take into account issues with branch lengths, but their simulations and our randomization tests show that they appear not to have been sufficient to correct for this bias (Figs. 2A and 2C). Dunn et al. also provided the hutan::picx() R function to compute PIC for OU trees. In their simulation-based function, they estimated ancestral states by the ‘GLS_OUS’ method using the bias calibrated phylogeny. Therefore, their method does not add anything specific to deal with the OU trees. Since they did not control for phylogenetic independence of the contrasts, and did not consider the relative ages of the speciation and old duplication events, they always obtained lower PIC of duplication events. Due to such phylogenetic internal parameter dependence, their PIC analyses produced similar trends with real or randomized data.

Assumptions of proper branch length information and of Brownian motion of trait evolution are related, so that modifications of branch lengths can change the evolutionary model (Diaz-Uriarte and Garland 1996; Díaz-Uriarte and Garland 1998). Contrasting a single rate OU to BM models, Dunn et al. (2018) identified 99.9% gene trees which favored an OU model, more explicitly an OU1 model. This appears to be 67% when we performed multivariate data modeling in a maximum likelihood framework on trees with less or no calibration bias (Supplementary fig. S8). PIC analyses with diagnostic tests provided weak support for the ortholog conjecture for the young duplicates (Figs. 3A–3C), in contrast to previous results of Dunn et al. Small effect size difference in our inference is not surprising since PIC is applied on OU trees. Similar patterns of results from empirical and randomization tests for the old duplicates indicate that one should be extremely careful before integrating them into a phylogenetic analysis. Branch length transformation attempts to transform the OU trees to BM trees to meet the underlying assumption of phylogenetic comparative method (Butler and King 2004). Hence, it can address the issue of low power when underlying assumptions of phylogenetic methods are violated (Diaz-Uriarte and Garland 1996; Díaz-Uriarte and Garland 1998). Following this approach along with the diagnostic tests, we obtained substantial support for the ortholog conjecture (Figs. 4A–4C, Supplementary figs. S9 and S10).

Phylogenetic data modeling also appears to be a powerful tool for such hypothesis testing, where one can estimate the trait evolutionary rates or optima shift rates per event without transforming OU trees to BM trees. More support for the OU trees (Supplementary fig. S8) could be due to the fact that we performed multivariate evolutionary model-fitting mostly on small trees (Cooper et al. 2016b). Among them only 8.6% trees supported the OUM model. Following the recommendation of Cooper et al. (2016), we applied Bayesian approach on small trees to accurately identify multi optima trees. Although previous studies (Uyeda and Harmon 2014; Khabbazian et al. 2016; Uyeda et al. 2017) have suggested a liberal cutoff of ≥ 0.2 to detect an optimum shift with a Bayesian approach, we used a strict posterior probability cutoff of ≥ 0.7. We performed our analyses on the 33.9% OUM trees passing such a strict posterior probability threshold. Our results from the PIC analyses with controls was also supported by the maximum likelihood, and Bayesian data modeling approaches. This shows that once proper precautions are taken, the empirical trends do not depend on the number of selected gene trees or of internal node events included.

Empirical support for the ortholog conjecture has been mixed, with some studies supporting it (Koonin 2005; Studer and Robinson-Rechavi 2009; Altenhoff et al. 2012; Chen and Zhang 2012; Gabaldón and Koonin 2013; Rogozin et al. 2014; Kryuchkova-Mostacci and Robinson-Rechavi 2016; Fukushima and Pollock 2020), and a few failing to do so (Nehrt et al. 2011; Dunn et al. 2018; Stamboulian et al. 2020). Our results provide additional support for the ortholog conjecture using tissue specificity data in a phylogenetic framework after controlling for biases. Due to lack of detailed functional information, many studies are still limited to gene expression data as a proxy of function. Recently, using functional replaceability assay, experimental studies (Kachroo et al. 2015; Laurent et al. 2020) have shown that orthologous genes can be swapped between essential yeast genes and human, although this is rarely the case for all the members of expanded human gene families (Laurent et al. 2020), validating one prediction of the ortholog conjecture.

## Materials and Methods

### Data reproducibility details

Our analyses are based on 21124 gene trees obtained from ENSEMBL Compara v.75 (Herrero et al. 2016) as used by Dunn et al. (2018). We used the same random seed number as in Dunn et al. (2018) to reproduce the simulation results for reanalysis. All reproduced data of Dunn et al. were stored in the “manuscript_dunn.RData” file (https://doi.org/10.5281/zenodo.4003391). We used the results stored in the ‘data’ or ‘phylo’ slot of the trees for further analyses. To differentiate our own function from theirs (Dunn et al. 2018), we renamed the original function script of Dunn et al. from “functions.R” to “functions_Dunn.R”. We made separate scripts for PIC analyses (“Premanuscript_run_TMRR.R”), and for data modeling analyses (“Model_fitting.R”). Some of the analyses were time consuming, so we stored our outputs in “Analyses_TMRR.RData”, and in “Model_fitting_TMRR.Rdata” files (https://doi.org/10.5281/zenodo.4003391), to load during analyses. All the details of different functions are provided inside the scripts. We supply all the previously stored data (to reduce computation time during reproduction of result) and function files including our own (“functions_TM_new.R”) with this manuscript. All scripts are available on GitHub: https://github.com/tbegum/Testing_the_ortholog_conjecture.

### Fixing time calibration bias of duplication nodes

We first present the approach that Dunn et al. (2018) used, for clarity. When two speciation nodes had the same label in the gene tree, Dunn et al. edited the more recent one to “NA” rather than “speciation”. Indeed the presence of the same clade names at different node depths forces all the intervening branches to have length zero when the tree is time calibrated, leading to failure of calibration (Dunn et al. 2018). For trait evolution, they annotated the tips of these modified trees with precomputed tissue specificity data, *τ* from 8 vertebrate species (human, gorilla, chimpanzee, macaque, mouse, opossum, platypus, and chicken) (from Kryuchkova-Mostacci and Robinson-Rechavi 2016). *τ* is a univariate index between 0 and 1 that measures tissue-specificity of gene expression (Yanai et al. 2005): *τ* close to 1 indicates high tissue specificity, while close to 0 indicates more ubiquitous expression. Here *τ* was computed across 6 tissues: brain, cerebellum, heart, kidney, liver, and testis, based on the RNA-seq data of Brawand et al. (2011). Dunn et al. pruned the gene trees to remove tips with missing *τ* data, and then time calibrated them using speciation clade ages in the chronos() function with the ‘correlated’ model from the R package “ape” (Paradis et al. 2004). The modified NA clades were not used for this calibration. They used 7 speciation time points with a maximum age of 296 My. Thus they obtained 8520 calibrated gene trees having at least 4 tips with non-null trait data (Table 1; Supplementary figs. S4A-S4C). Among these trees, 2990 were pure speciation trees, which includes 12919 speciation events, or 19% of all speciation nodes.

Relative to Dunn et al., we exchanged the order of pruning and time calibration steps, i.e., we first time calibrated the 21124 modified (i.e. with NA added) gene trees, followed by pruning to have at least 4 tips with *τ* data. This makes use of all 32 available speciations time points, and helps to limit the calibration bias of the old duplication events (Supplementary figs. S4D-S4F). Calibration fails for some trees, and we obtained 7336 calibrated gene trees. The maximum node age of old duplication events is 1175.2 My for these trees, as opposed to 11799977 My (older than the universe) for the trees obtained by the original approach (Table 1, Supplementary table S1). Among these 7336 gene trees, we kept 4288 which have at least 1 speciation and 1 duplication events; we removed 39 pure duplication and 3009 pure speciation trees. This 4288 gene tree set is our basis for evaluating phylogenetic methods’ capacity to test the ortholog conjecture (Table 1): we compare the evolutionary rates, *σ*^2^, or PICs of speciation and duplication events of the same genes.

### Model selection for *τ* evolution

We followed a state dependent model-fitting approach to identify Brownian motion (BM) or Ornstein-Uhlenbeck (OU) trees. We classified time-calibrated gene duplication nodes as “young” (≤ 296 My, the maximum speciation age) or “old” (> 296 My) before model fitting. We performed stochastic mapping of our gene trees by assigning discrete states (“speciation”, “young-duplication”, “old-duplication”, and “NA”) to the branches based on the corresponding ancestral node events using the simmap() function of the phytools R package (Revell 2012). For each mapped tree, we fitted 4 different models of *τ* evolution using maximum-likelihood: (i) BM1, a single Brownian motion rate of evolution (i.e. σ^2^_speciation_ = σ^2^_young-duplication_ = σ^2^_old-duplication_), (ii) BMM, a BM with multiple rates of evolution for different events (i.e. different σ^2^ are allowed), (iii) OU1, a single optimum OU model (i.e. θ_speciation_ = θ_young-duplication_ θ_old-duplication_, σ^2^_speciation_ = σ^2^_young-duplication_ = σ^2^_old-duplication_, α_speciation_ = α_young-duplication_ = α_old-duplication_), and (iv) OUM, a multi optimum OU model with identical strength of selection and rate of drift acting on all selective regimes (i.e. like OU1 but θ_speciation_ ≠ θ_young-duplication_ ≠ θ_old-duplication_).

We used both the mvMORPH (Clavel et al. 2015), and OUwie (Beaulieu et al. 2012) R packages to perform model-fitting. Sometimes the information contained within a tree is insufficient with respect to the complexity of the fitted models. This can lead to poor model choice by returning a log-likelihood that is suboptimal and may provide incorrect estimation of one or more model parameters for that tree (Beaulieu et al. 2012). Hence, we included the diagnostics (diagnostic=T or diagn=T) during model-fitting. The eigen values of the Hessian matrix of the diagnostics indicate whether convergence of the model has been achieved or whether the parameter estimates are reliable (Beaulieu et al. 2012). For the BM1, BMM, OU1, and OUM models, we first fitted the model using mvMORPH for each gene tree. If any of the model failed to converge for the tree or if the eigen values of the Hessian matrix indicated that it was not reliable, we re-fitted that model using OUwie to include it in model comparison. If still it failed, we removed that model for that tree. For model comparisons on each gene tree, we calculated the Akaike weights (ω) for each fitted model by means of the second order Akaike information criteria (AICc), which includes a correction for small sample sizes (Akaike 1974; Burnham and Anderson 2002). The model with highest ω was selected as the best-supported model of *τ* evolution for the tree (Burnham and Anderson 2002; Gearty et al. 2018). We estimated model parameters for each tree based on the best fit model.

### Bayesian modeling to detect phenotypic optimum shift

Regime shifts, i.e. shifts of optimal *τ* values, in OU models were detected by a Bayesian phylogenetic approach of the bayou R package (Uyeda and Harmon 2014). The reversible-jump phylogenetic comparative approach was used to perform MCMC sampling of locations, magnitudes and numbers of shifts in multiple-optima Ornstein-Uhlenbeck models. We ran MCMC chains for 100000 generations, and the first 30% of samples were dropped as burn-in. We used a strict threshold of posterior probability ≥ 0.7 to detect an adaptive shift at a given branch of the phylogeny. For each event (“speciation” or “duplication”), we used a ratio of the number of optimum shifts to the number of branches for that event to estimate the proportions of shifts in a phylogeny.

### Randomization test of *τ* values

For each tree, we used *τ* data (column name “Tau” in each tree ‘data’ object) across the tips to carry out our randomization test. To randomize we permuted the actual *τ* data without altering internal node events. The pic() function of the “ape” package (Paradis et al. 2004) was used to compute PIC of nodes for each tree using permuted *τ* of tips. For each run, we compared the contrasts of speciation and duplication events of the whole set of randomized trees to estimate difference in event contrasts based on Wilcoxon signed rank test. For 100 runs, we repeated the above process 100 times to obtain a distribution plot of 100 independent *P* values. For our model-fitting approach, we used the same empirical simmap trees with permuted *τ* data at the tips. We re-estimated the model parameters of the randomized *τ* trees using the best fit model chosen for the corresponding empirical gene trees.

### Randomization test of node events

Some of the speciation nodes had daughters with same clade names in the gene trees we used for our study. Dunn et al. changed such node events to “NA” to avoid problems during time calibration of the trees. Such annotated node event information (“Speciation”, “Duplication”, “NA”) for each tree was available as “Event” in the tree ‘data’ slot. To randomize, we permuted the internal node events (added as column name “event_new” in the ‘data’ slot) by maintaining the actual proportion of events for each tree. Then, we used the PIC of actual *τ* at tips to estimate contrasts difference between newly assigned speciation and duplication node events by Wilcoxon rank tests. For 100 independent runs, we repeated the same procedure to obtain 100 independent *P* values. Since the internal node events were changed after such randomization, we reclassified gene duplication nodes as “young” or “old” on the event modified trees, and repainted the trees. We re-estimated the model parameters for the discrete states of the randomized events trees using the best fit model chosen for the corresponding empirical gene trees.

### Checking for contrasts standardization by diagnostic tests

We used several additional diagnostic tests on those trees to identify adequate independent nodes contrast standardization before drawing any inference by PIC method, as recommended in several studies (Garland 1992; Diaz-Uriarte and Garland 1996; Díaz-Uriarte and Garland 1998; Freckleton and Harvey 2006; Cooper et al. 2016a). The most usual method for contrasts standardization is to check a correlation between the absolute values of PICs and their expected standard deviations (i.e. square root of sum of branch lengths) (Garland et al. 1992; Díaz-Uriarte and Garland 1998; Cooper et al. 2016a). Under Brownian motion, there should be no correlation. This test and the correlation between the absolute values of PICs and the logarithm of their node age are model diagnostic plot tests in the caper (“Comparative Analyses of Phylogenetics and Evolution in R”) package (Purvis and Rambaut 1995; Cooper et al. 2016a; Orme 2018; R Core Team 2018). We used both of them by using the “crunch” algorithm of the caper package, which implements the methods originally provided in CAIC (Purvis and Rambaut 1995; Cooper et al. 2016a; Orme 2018; R Core Team 2018). Correlation of node heights with absolute values of contrasts or PICs has also been reported to be a reliable indicator of deviation from the Brownian model (Freckleton and Harvey 2006). Hence, we computed node height for each node in a tree using the ape package (Paradis et al. 2004). We also used the correlations of node height and node depth to the absolute value of nodes contrasts to rule out significant trend in any of the 4 tests. We used *P* < 0.05 to assess a significant correlation for the diagnostic tests. A significant trend (positive or negative) indicates phylogenetic dependence for that tree (Garland 1992; Garland et al. 1992; Díaz-Uriarte and Garland 1998; Freckleton and Harvey 2006; Cooper et al. 2016a), and we removed those trees from our analysis. Contrast calculation on negative branch lengths is not desirable, so we removed trees with negative branch lengths before applying the crunch() function. To assure that nodes contrast standardization is independent of the phylogeny, we considered sets of trees passing all 4 diagnostic tests for further analyses.

### Branch length transformation

Transformation of branch lengths has been proposed to restore the performance of PIC method when the true evolutionary model is not BM or is unknown, or when branch lengths are in error (Garland et al. 1992; Diaz-Uriarte and Garland 1996; Díaz-Uriarte and Garland 1998). In such cases, branch lengths are transformed by raising a family power of branch length ranging from 0 to 2 in intervals of 0.1, plus the log10 of the branch lengths (Diaz-Uriarte and Garland 1996; Díaz-Uriarte and Garland 1998). For each transformation, the program computes the correlation between the absolute value of the standardized contrasts and their standard deviations until no significant correlation is obtained, to ensure adequate independent contrasts standardization (Diaz-Uriarte and Garland 1996; Díaz-Uriarte and Garland 1998). Finally, we excluded trees for which adequate contrasts standardization is not achieved even after raising the branch length power to 2 (Diaz-Uriarte and Garland 1996; Díaz-Uriarte and Garland 1998).

### Details of other packages used in this study

We used phylosig function() of the phytools package (Revell 2012) to identify trees with phylogenetic signal (*P* < 0.05) using Blomberg’s K (Blomberg et al. 2003; Münkemüller et al. 2012; Revell 2012). Analyses and plotting were performed in R version 3.5.1 (R Core Team 2018) using treeio (Guangchuang 2018), ggtree (Guangchuang et al. 2017), stringr (Wickham 2019), digest (Antoine Lucas et al. 2018), dplyr (Wickham et al. 2017), tidyverse (Wickham 2017), ggrepel (Slowikowski 2018), gtools (Warnes et al. 2018), ggplot2 (Wickham 2016), cowplot (Wilke 2019), easyGgplot2 (Kassambara 2014), gridExtra (Auguie 2017), and png (Urbanek 2013) libraries.

## Acknowledgements

We sincerely acknowledge Martha Liliano Serranno Serranno for initial help with the phylogenetic independent contrast method. We thank Nicolas Salamin, Julien Wollbrett, Jialin Liu, Sebastien Moretti, Sara Fonseca Costa, Kamil Jaron and all the members of the Robinson-Rechavi group for their help and useful discussions. We also acknowledge Cassey Dunn for his initial help in reproducing their results. Parts of the computations were performed at the Vital-IT (http://www.vital-it.ch) Center for high-performance computing of the SIB Swiss Institute of Bioinformatics.

## Supporting Information

**Figure S1: Expectations from phylogenetic and pairwise comparison approaches under null and ortholog conjecture scenarios.** PIC: Phylogenetic Independent Contrast, OC: Ortholog Conjecture. We present 4 time-calibrated gene trees of Dunn et al. (2018) as illustration. Trees A and B are well calibrated, with the duplication ages are constrained by speciation ages, as shown by the time scales below each phylogeny. Trees C and D represent biased calibrated trees, where old duplication branches are inaccurately calibrated due to lack of age constraints. To evaluate the impacts of gene duplication and speciation events in trait evolution, pairwise comparisons do not rely on the branch lengths of a calibrated phylogeny, but phylogenetic methods do. If time calibration of old duplication nodes has no influence in the inference of phylogenetic approaches, we expect to obtain patterns under a null and OC scenarios as shown in the right part of the figure. This means that the phylogenetic contrasts or pairwise correlations of different events should be drawn from the same distribution under a null model, while the expectation differs under the OC model. We used 2 times higher rates of trait evolution (*τ* here) following duplications than speciations (i.e. σ^2^_duplication_ = 2 * σ^2^_speciation_) in this example for the OC model.

**Figure S2.: Repeating simulations on all calibrated trees with different random seed number.** *P* values are from Wilcoxon two-tailed tests. Simulations with different seed number did not change the trend of results as reported in Fig. 1A.

**Figure S3: Simulation analyses on 1135 trees with strong phylogenetic signals.** *P* values are from Wilcoxon two-tailed tests. Dunn et al. used a cutoff of K > 0.551 to identify trees with strong phylogenetic signals. However, trees with higher K statistic can have corresponding *P* values which are non-significant. Considering both K statistic and *P* value, we found similar trends as was observed with 2082 trees.

**Figure S4: Difference between time calibration approaches of Dunn et al. (2018) and of this study.** In this example, we used the phylogeny of ACP1 gene. The top panel (A-C) shows the steps used by Dunn et al. (2018), while the bottom panel (D-F) shows the steps used in this study. Gene trees obtained from Ensembl (Herrero et al. 2016) have branch lengths in substitutions per site. (A) and (D) are the same gene tree, where Dunn et al. (2018) edited few speciation events to ‘NA’ to pass the time calibration step. (B) The gene trees are pruned to species with available *τ*. (C) The pruned tree is time calibrated using speciation time points. Pruning before time calibration produces tree with many duplications, and NA nodes older to the oldest speciation nodes as in (B). This leads to using only 7 speciation time points for calibration. Due to unavailable age constraints on the old duplication nodes, the time scale of the phylogeny in (C) reaches 880 million years (My). When we performed time calibration before pruning as in (E), we could use 32 speciation nodes for time calibration. This means that we could use many speciation nodes for time calibration, although τ data was unavailable for species at tips due to the choice of species in this study. Hence, the old duplication nodes are constrained by the age of speciation nodes older to them, and thus the maximum age is now of 356 My (F).

**Figure S5: Re-analyses of expected variances of calibrated trees considered by Dunn et al.** The expected variance plots of (A) all 8520 calibrated trees, and (B) 2082 trees with strong phylogenetic signal. The dotted line represents the mean expected variance of the events. These plots show why duplication nodes preceding ancient speciation nodes can be problematic for PIC.

**Figure S6: *P* value distribution plots after 100 independent runs on each set of trees.** Wilcoxon two-tailed test with 95% confidence interval was used to compare the speciation and duplication contrasts after randomization tests. (A) and (B) applied to trees with at least one speciation and one duplication event. (C) and (D) applied to trees with strong phylogenetic signal. (A) and (C) randomization of trait (τ) over the trees. (B) and (D) randomization of internal node events. The inset plots show *P* values adjusted with Benjamini-Hochberg (Benjamini and Yekutieli 2005; Hochberg and Benjamini 1990). Supporting our observations of Figs. 2A and 2C, all the plots confirm that the empirical result of Dunn et al. (2018) is not different from randomized test results.

**Figure S7: The ortholog conjecture test after randomizations of contrasts standardized trees.** *P* values are from Wilcoxon two-tailed tests. ‘PICs’: Phylogenetic Independent Contrasts. Values inside boxplots denote median PIC value of the corresponding event. (A-B) Plots after randomizing τ, and after randomizing events using the same trees as in Fig. 3.

**Figure S8: Multivariate model fitting result using a maximum likelihood framework.** BM1: Single rate Brownian; BMM: Multi rates Brownian; OU1: Single optimum Ornstein-Uhlenbeck; and OUM: Multi optima Ornstein-Uhlenbeck models.

**Figure S9: The ortholog conjecture test for τ on calibrated trees of Dunn et al.** *P* values are from Wilcoxon two-tailed tests. ‘PICs’: Phylogenetic Independent Contrasts. Values inside boxplots denote median PIC value of the corresponding event. (A) Using 8417 out of 8520 calibrated trees that passed diagnostic tests following branch length transformation. (B) Plot after randomizing τ, and (C) after randomizing events using the same branch transformed trees as in (A).

**Figure S10: The ortholog conjecture test for τ on branch transformed trees with strong phylogenetic signals.** *P* value are from Wilcoxon two-tailed tests. ‘PICs’: Phylogenetic Independent Contrasts. Values inside boxplots denote median PIC value of the corresponding event. (A) using 2080 out of 2082 calibrated trees that passed diagnostic tests following branch length transformation. (B) Plot after randomizing τ, and (C) after randomizing events using the same branch transformed trees as in (A).

**Table S1:** Summary statistics of calibrated old duplication nodes for 8420 trees of Dunn et al. (2018).

| **Maximum age group in**<br>**Million Years (My)** | **Count** |
| --- | --- |
| 296-500 | 2917 |
| 501-900 | 7548 |
| 901-3000 | 49 |
| 3001-10000 | 16 |
| 10001-11799977 | 9 |

**Table S2:** Analyses on multi optima OU trees.

| **Duplication**<br>**age** | **Data** | $\theta_{\text{Speciation}}$ | $\theta_{\text{Duplication}}$ | $\theta_{\text{Duplication}} / \theta_{\text{Speciation}}$ | ***P*-value** |
| --- | --- | --- | --- | --- | --- |
| Young | Empirical<br>(nSpeciation= 2690;<br>nDuplication = 842) | 0.41 | 0.74 | 1.8 | $8.6\text{e}^{-10}$ |
| | Randomized $\tau$<br>(nSpeciation= 2690;<br>nDuplication = 842) | 0.53 | 0.55 | 1.03 | 0.97 |
|  | Randomized<br>events<br>(nSpeciation= 1872;<br>nDuplication = 698) | 0.50 | 0.53 | 1.06 | 0.75 |
| Old | Empirical<br>(nSpeciation= 4152;<br>nDuplication = 847) | 0.42 | 0.92 | 2.19 | $2.4\text{e}^{-4}$ |
| | Randomized $\tau$<br>(nSpeciation= 4152;<br>nDuplication = 847) | 0.54 | $1.5\text{e}^{-11}$ | $2.8\text{e}^{-11}$ | $1.3\text{e}^{-07}$ |
|  | Randomized<br>events<br>(nSpeciation= 2081;<br>nDuplication = 482) | 0.51 | 0.66 | 1.29 | 0.73 |

964 *Note:* Median values of $\sigma^2$ are shown. *P*-value from paired two-sided Wilcoxon test.

**Table S3:** Summary statistics on OUM trees, passing both the maximum likelihood and the Bayesian approaches with a posterior probability cutoff of ≥ 0.7.

| Duplication<br>Age | Proportions of regime shifts per branch |  | Paired two-sided<br>Wilcoxon rank sum test | Regime shift rates (shifts/My) |  | Two-sided<br>Wilcoxon rank test |
| --- | --- | --- | --- | --- | --- | --- |
|  | After speciation | After duplication |  | After speciation | After duplication |  |
| Young | 2.8% | 8.3% | $8.3e^{-4}$ | 0.012 | 0.032 | $6.7e^{-6}$ |
| Old | 0% | 16.7% | $< 2.2e^{-16}$ | 0.012 | 0.0025 | $< 2.2e^{-16}$ |

*Note-* Above analyses include 2779 speciation, 486 young, and 548 old duplication events.
Values shown in the table indicate median values. The difference in proportions of regime shifts per branch after speciation events for two types of duplications is due to the different sets of trees used. Few trees shared both types of duplicates. Proportions of regime shifts per branch of events is estimated for each tree, and thus paired Wilcoxon test is used to compare the difference. A single gene tree can have one or many optima shift rate(s) for events, and thus two-sided Wilcoxon rank test was used for comparison.

## References

Akaike H. 1974. New look at statistical-model identification. Automatic Control, IEEE Transactions on 19:716–723.

Altenhoff AM, Studer RA, Robinson-Rechavi M, Dessimoz C. 2012. Resolving the ortholog conjecture: Orthologs tend to be weakly, but significantly, more similar in function than paralogs. PLoS Comput Biol 8:e1002514. Available from: https://www.ncbi.nlm.nih.gov/pubmed/22615551

Antoine Lucas DE with contributions by, Tuszynski J, Bengtsson H, Urbanek S, Frasca M, Lewis B, Stokely M, Muehleisen H, Murdoch D, Hester J, et al. 2018. Digest: Create compact hash digests of r objects. Available from: https://CRAN.R-project.org/package=digest

Auguie B. 2017. GridExtra: Miscellaneous functions for “grid” graphics. Available from: https://CRAN.R-project.org/package=gridExtra

Beaulieu JM, Jhwueng DC, Boettiger C, O’Meara BC. 2012. Modeling stabilizing selection: Expanding the Ornstein-Uhlenbeck model of adaptive evolution. Evolution 66:2369–2383. Available from: https://www.ncbi.nlm.nih.gov/pubmed/22834738

Benjamini Y, Yekutieli D. 2005. Quantitative trait loci analysis using the false discovery rate. Genetics 171:783–790. Available from: https://www.ncbi.nlm.nih.gov/pubmed/15956674

Blomberg SP, Garland T, Ives AR. 2003. Testing for phylogenetic signal in comparative data: Behavioral traits are more labile. Evolution 57:717–745. Available from: https://www.ncbi.nlm.nih.gov/pubmed/12778543

Brawand D, Soumillon M, Necsulea A, Julien P, Csárdi G, Harrigan P, Weier M, Liechti A, Aximu-Petri A, Kircher M, et al. 2011. The evolution of gene expression levels in mammalian organs. Nature 478:343–348. Available from: https://www.ncbi.nlm.nih.gov/pubmed/22012392

Burnham K, Anderson D. 2002. Model selection and multimodel inference. In: Springer, New York,

Butler MA, King AA. 2004. Phylogenetic comparative analysis: a modeling approach for adaptive evolution. Am Nat 164:683–695. Available from: https://www.ncbi.nlm.nih.gov/pubmed/29641928

Catalán A, Briscoe AD, Höhna, S. 2019. Drift and directional selection are the evolutionary forces driving gene expression divergence in eye and brain tissue of. Genetics 213:581–594. Available from: https://www.ncbi.nlm.nih.gov/pubmed/31467133

Chen J, Swofford R, Johnson J, Cummings BB, Rogel N, Lindblad-Toh K, Haerty W, Palma FD, Regev A. 2019. A quantitative framework for characterizing the evolutionary history of mammalian gene expression. Genome Res 29:53–63. Available from: https://www.ncbi.nlm.nih.gov/pubmed/30552105

Chen X, Zhang J. 2012. The ortholog conjecture is untestable by the current gene ontology but is supported by rna sequencing data. PLoS Comput Biol 8:e1002784. Available from: https://www.ncbi.nlm.nih.gov/pubmed/23209392

Clavel J, Escarguel G, Merceron G. 2015. mvMORPH: An R package for fitting multivariate evolutionary models to morphometric data. Methods Ecol Evol 6:1311–1319.

Conant GC, Wagner A. 2003. Asymmetric sequence divergence of duplicate genes. Genome Res 13:2052–2058. Available from: https://www.ncbi.nlm.nih.gov/pubmed/12952876

Cooper N, Thomas GH, FitzJohn RG. 2016a. Shedding light on the ‘dark side’ of phylogenetic comparative methods. Methods Ecol Evol 7:693–699. Available from: https://www.ncbi.nlm.nih.gov/pubmed/27499839

Cooper N, Thomas GH, Venditti C, Meade A, Freckleton RP. 2016b. A cautionary note on the use of ornstein uhlenbeck models in macroevolutionary studies. Biol J Linn Soc Lond 118:64–77. Available from: https://www.ncbi.nlm.nih.gov/pubmed/27478249

Cornwell W, Nakagawa S. 2017. Phylogenetic comparative methods. Curr Biol 27:R333–R336. Available from: https://www.ncbi.nlm.nih.gov/pubmed/28486113

Diaz-Uriarte R, Garland T. 1996. Testing hypotheses of correlated evolution using phylogenetically independent contrasts: Sensitivity to deviations from brownian motion. Syst Biol 45:27–47.

Díaz-Uriarte R, Garland T. 1998. Effects of branch length errors on the performance of phylogenetically independent contrasts. Syst Biol 47:654–672. Available from: https://www.ncbi.nlm.nih.gov/pubmed/12066309

Dunn CW, Zapata F, Munro C, Siebert S, Hejnol A. 2018. Pairwise comparisons across species are problematic when analyzing functional genomic data. Proc Natl Acad Sci U S A 115:E409–E417. Available from: https://www.ncbi.nlm.nih.gov/pubmed/29301966

Eastman JM, Alfaro ME, Joyce P, Hipp AL, Harmon LJ. 2011. A novel comparative method for identifying shifts in the rate of character evolution on trees. Evolution 65:3578–3589. Available from: https://www.ncbi.nlm.nih.gov/pubmed/22133227

Eng KH, Bravo HC, Keleş S. 2009. A phylogenetic mixture model for the evolution of gene expression. Mol Biol Evol 26:2363–2372. Available from: https://www.ncbi.nlm.nih.gov/pubmed/19602540

Felsenstein J. 1985. Phylogenies and the comparative method. Am Nat 125:1–15. Available from: https://www.jstor.org/stable/2461605

Freckleton R. 2000. Phylogenetic tests of ecological and evolutionary hypotheses: Checking for phylogenetic independence. Func Ecol 14:129–134.

Freckleton R, Harvey P, Pagel M. 2002. Phylogenetic analysis and comparative data: A test and review of evidence. Am Nat 160:712–726.

Freckleton RP, Harvey PH. 2006. Detecting non-brownian trait evolution in adaptive radiations. PLoS Biol 4:e373. Available from: https://www.ncbi.nlm.nih.gov/pubmed/17090217

Fukushima K, Pollock DD. 2020. Organ-specific propensity drives patterns of gene expression evolution. BioRxiv. doi: https://doi.org/10.1101/409888

Gabaldón T, Koonin EV. 2013. Functional and evolutionary implications of gene orthology. Nat Rev Genet 14:360–366. Available from: https://www.ncbi.nlm.nih.gov/pubmed/23552219

Garland T. 1992. Rate tests for phenotypic evolution using phylogenetically independent contrasts. Am Nat 140:509–519. Available from: https://www.ncbi.nlm.nih.gov/pubmed/19426053

Garland TJ, Harvey P, Ives A. 1992. Procedure for the analysis of comparative data using phylogenetically independent contrasts. Syst Biol 41:18–32.

Gearty W, McClain CR, Payne JL. 2018. Energetic tradeoffs control the size distribution of aquatic mammals. Proc Natl Acad Sci USA 115:4194–4199. Available from: https://www.ncbi.nlm.nih.gov/pubmed/29581289

Grafen A. 1989. The phylogenetic regression. Philos Trans R Soc Lond B Biol Sci 326:119–157. Available from: https://www.ncbi.nlm.nih.gov/pubmed/2575770

Guangchuang Y. 2018. Treeio: Base classes and functions for phylogenetic tree input and output. Available from: https://guangchuangyu.github.io/software/treeio

Guangchuang Y, David S, Huachen Z, Yi G, Tommy T-YL. 2017. Ggtree: An R package for visualization and annotation of phylogenetic trees with their covariates and other associated data. Methods Ecol Evol 8:28–36.

Han MV, Demuth JP, McGrath CL, Casola C, Hahn MW. 2009. Adaptive evolution of young gene duplicates in mammals. Genome Res 19:859–867. Available from: https://www.ncbi.nlm.nih.gov/pubmed/19411603

Hansen TF. 1997. Stabilizing selection and the comparative analysis of adaptation. Evolution 51:1341–1351. Available from: https://www.ncbi.nlm.nih.gov/pubmed/28568616

Herrero J, Muffato M, Beal K, Fitzgerald S, Gordon L, Pignatelli M, Vilella AJ, Searle SM, Amode R, Brent S, et al. 2016. Ensembl comparative genomics resources. Database. Available from: https://www.ncbi.nlm.nih.gov/pubmed/27141089

Hochberg Y, Benjamini Y. 1990. More powerful procedures for multiple significance testing. Stat Med 9:811–818. Available from: https://www.ncbi.nlm.nih.gov/pubmed/2218183

Holland PW, Marlétaz F, Maeso I, Dunwell TL, Paps J. 2017. New genes from old: Asymmetric divergence of gene duplicates and the evolution of development. Philos Trans R Soc Lond B Biol Sci 372. Available from: https://www.ncbi.nlm.nih.gov/pubmed/27994121

Kachroo AH, Laurent JM, Yellman CM, Meyer AG, Wilke CO, Marcotte EM. 2015. Evolution. systematic humanization of yeast genes reveals conserved functions and genetic modularity. Science 348:921–925. Available from: https://www.ncbi.nlm.nih.gov/pubmed/25999509

Kassambara A. 2014. EasyGgplot2: Perform and customize easily a plot with ggplot2. Available from: http://www.sthda.com

Khabbazian M, Kriebel R, Rohe K, Ané C. 2016. Fast and accurate detection of evolutionary shifts in ornstein-uhlenbeck models. Methods Ecol Evol 7:811–824.

Kim SH, Yi SV. 2006. Correlated asymmetry of sequence and functional divergence between duplicate proteins of saccharomyces cerevisiae. Mol Biol Evol 23:1068–1075. Available from: https://www.ncbi.nlm.nih.gov/pubmed/16510556

Koonin EV. 2005. Orthologs, paralogs, and evolutionary genomics. Annu Rev Genet 39:309–338. Available from: https://www.ncbi.nlm.nih.gov/pubmed/16285863

Kryuchkova-Mostacci N, Robinson-Rechavi M. 2016. Tissue-specificity of gene expression diverges slowly between orthologs, and rapidly between paralogs. PLoS Comput Biol 12:e1005274. Available from: https://www.ncbi.nlm.nih.gov/pubmed/28030541

Laurent JM, Garge RK, Teufel AI, Wilke CO, Kachroo AH, Marcotte EM. 2020. Humanization of yeast genes with multiple human orthologs reveals functional divergence between paralogs. PLoS Biol 18:e3000627. Available from: https://www.ncbi.nlm.nih.gov/pubmed/32421706

Martins E, Hansen T. 1997. Phylogenies and the comparative method: A general approach to incorporating phylogenetic information into the analysis of interspecific data. Am Nat 149:646–667.

Molina-Venegas R, Rodríguez M. 2017. Revisiting phylogenetic signal; strong or negligible impacts of polytomies and branch length information? BMC Evol Biol 17:53. Available from: https://www.ncbi.nlm.nih.gov/pubmed/28201989

Münkemüller T, Lavergne S, Bzeznik B, Dray S, Jombart T, Schiffers K, Thuiller W. 2012. How to measure and test phylogenetic signal. Methods Ecol Evol 3:743–756.

Nehrt NL, Clark WT, Radivojac P, Hahn MW. 2011. Testing the ortholog conjecture with comparative functional genomic data from mammals. PLoS Comput Biol 7:e1002073. Available from: https://www.ncbi.nlm.nih.gov/pubmed/21695233

Oakley TH, Gu Z, Abouheif E, Patel NH, Li WH. 2005. Comparative methods for the analysis of gene-expression evolution: An example using yeast functional genomic data. Mol Biol Evol 22:40–50. Available from: https://www.ncbi.nlm.nih.gov/pubmed/15356281

Oakley TH, Ostman B, Wilson AC. 2006. Repression and loss of gene expression outpaces activation and gain in recently duplicated fly genes. Proc Natl Acad Sci U S A 103:11637–11641. Available from: https://www.ncbi.nlm.nih.gov/pubmed/16864793

O’Meara BC, Ané C, Sanderson MJ, Wainwright PC. 2006. Testing for different rates of continuous trait evolution using likelihood. Evolution 60:922–933. Available from: https://www.ncbi.nlm.nih.gov/pubmed/16817533

Orme D. 2018. The caper package: Comparative analysis of phylogenetics and evolution in R. Available from: https://cran.r-project.org/web/packages/caper/vignettes/caper.pdf

Pagel M. 1999. Inferring the historical patterns of biological evolution. Nature 401:877–884.

Panchin AY, Gelfand MS, Ramensky VE, Artamonova II. 2010. Asymmetric and non-uniform evolution of recently duplicated human genes. Biol Direct 5:54. Available from: https://www.ncbi.nlm.nih.gov/pubmed/20825637

Paradis E, Claude J, Strimmer K. 2004. APE: Analyses of phylogenetics and evolution in R language. Bioinformatics 20:289–290. Available from: https://www.ncbi.nlm.nih.gov/pubmed/14734327

Pegueroles C, Laurie S, Albà MM. 2013. Accelerated evolution after gene duplication: A time-dependent process affecting just one copy. Mol Biol Evol 30:1830–1842. Available from: https://www.ncbi.nlm.nih.gov/pubmed/23625888

Pennell MW, Eastman JM, Slater GJ, Brown JW, Uyeda JC, FitzJohn RG, Alfaro ME, Harmon LJ. 2014. Geiger v2.0: An expanded suite of methods for fitting macroevolutionary models to phylogenetic trees. Bioinformatics 30:2216–2218. Available from: https://www.ncbi.nlm.nih.gov/pubmed/24728855

Pich I Roselló O, Kondrashov FA. 2014. Long-term asymmetrical acceleration of protein evolution after gene duplication. Genome Biol Evol 6:1949–1955. Available from: https://www.ncbi.nlm.nih.gov/pubmed/25070510

Purvis A, Rambaut A. 1995. Comparative analysis by independent contrasts (caic): An apple macintosh application for analysing comparative data. Comput Appl Biosci 11:247–251. Available from: https://www.ncbi.nlm.nih.gov/pubmed/7583692

R Core Team. 2018. R: A language and environment for statistical computing. Vienna, Austria: R Foundation for Statistical Computing Available from: https://www.R-project.org/

Revell LJ. 2012. Phytools: An R package for phylogenetic comparative biology (and other things). Methods Ecol Evol 3:217–223.

Rogozin IB, Managadze D, Shabalina SA, Koonin EV. 2014. Gene family level comparative analysis of gene expression in mammals validates the ortholog conjecture. Genome Biol Evol 6:754–762. Available from: https://www.ncbi.nlm.nih.gov/pubmed/24610837

Rohlf FJ. 2001. Comparative methods for the analysis of continuous variables: Geometric interpretations. Evolution 55:2143–2160. Available from: https://www.ncbi.nlm.nih.gov/pubmed/11794776

Rohlfs RV, Nielsen R. 2015. Phylogenetic ANOVA: The expression variance and evolution model for quantitative trait evolution. Syst Biol 64:695–708. Available from: https://www.ncbi.nlm.nih.gov/pubmed/26169525

Sanderson MJ. 2002. Estimating absolute rates of molecular evolution and divergence times: A penalized likelihood approach. Mol Biol Evol 19:101–109. Available from: https://www.ncbi.nlm.nih.gov/pubmed/11752195

Scannell DR, Wolfe KH. 2008. A burst of protein sequence evolution and a prolonged period of asymmetric evolution follow gene duplication in yeast. Genome Res 18:137–147. Available from: https://www.ncbi.nlm.nih.gov/pubmed/18025270

Slowikowski K. 2018. Ggrepel: Automatically position non-overlapping text labels with ‘ggplot2’. Available from: https://CRAN.R-project.org/package=ggrepel

Sonnhammer EL, Gabaldón T, Sousa da Silva AW, Martin M, Robinson-Rechavi M, Boeckmann B, Thomas PD, Dessimoz C, consortium Q for O. 2014. Big data and other challenges in the quest for orthologs. Bioinformatics 30:2993–2998. Available from: https://www.ncbi.nlm.nih.gov/pubmed/25064571

Stamboulian M, Guerrero RF, Hahn MW, Radivojac P. 2020. The ortholog conjecture revisited: The value of orthologs and paralogs in function prediction. Bioinformatics 36:i219–i226. Available from: https://www.ncbi.nlm.nih.gov/pubmed/32657391

Studer RA, Robinson-Rechavi M. 2009. How confident can we be that orthologs are similar, but paralogs differ? Trends Genet 25:210–216. Available from: https://www.ncbi.nlm.nih.gov/pubmed/19368988

Thomas GH, Freckleton RP, Székely T. 2006. Comparative analyses of the influence of developmental mode on phenotypic diversification rates in shorebirds. Proc Biol Sci 273:1619–1624. Available from: https://www.ncbi.nlm.nih.gov/pubmed/16769632

Urbanek S. 2013. Png: Read and write png images. Available from: https://CRAN.R-project.org/package=png

Uyeda JC, Harmon LJ. 2014. A novel bayesian method for inferring and interpreting the dynamics of adaptive landscapes from phylogenetic comparative data. Syst Biol 63:902–918. Available from: https://www.ncbi.nlm.nih.gov/pubmed/25077513

Uyeda JC, Pennell MW, Miller ET, Maia R, McClain CR. 2017. The evolution of energetic scaling across the vertebrate tree of life. Am Nat 190:185–199. Available from: https://www.ncbi.nlm.nih.gov/pubmed/28731792

Warnes GR, Bolker B, Lumley T. 2018. Gtools: Various R programming tools. Available from: https://CRAN.R-project.org/package=gtools

Wickham H. 2016. Ggplot2: Elegant graphics for data analysis. Springer-Verlag New York Available from: https://ggplot2.tidyverse.org

Wickham H. 2017. Tidyverse: Easily install and load the ‘tidyverse’. Available from: https://CRAN.R-project.org/package=tidyverse

Wickham H. 2019. Stringr: Simple, consistent wrappers for common string operations. Available from: https://CRAN.R-project.org/package=stringr

Wickham H, Francois R, Henry L, Müller K. 2017. dplyr: A grammar of data manipulation. Available from: https://CRAN.R-project.org/package=dplyr

Wilke CO. 2019. Cowplot: Streamlined plot theme and plot annotations for ‘ggplot2’. Available from: https://CRAN.R-project.org/package=cowplot

Yanai I, Benjamin H, Shmoish M, Chalifa-Caspi V, Shklar M, Ophir R, Bar-Even A, Horn-Saban S, Safran M, Domany E, et al. 2005. Genome-wide midrange transcription profiles reveal expression level relationships in human tissue specification. Bioinformatics 21:650–659. Available from: <Go to ISI>://WOS:000227241200012

