## Supplementary figures and images for "Special care is needed in applying phylogenetic comparative methods to gene trees with speciation and duplication nodes"

### supplementary figures

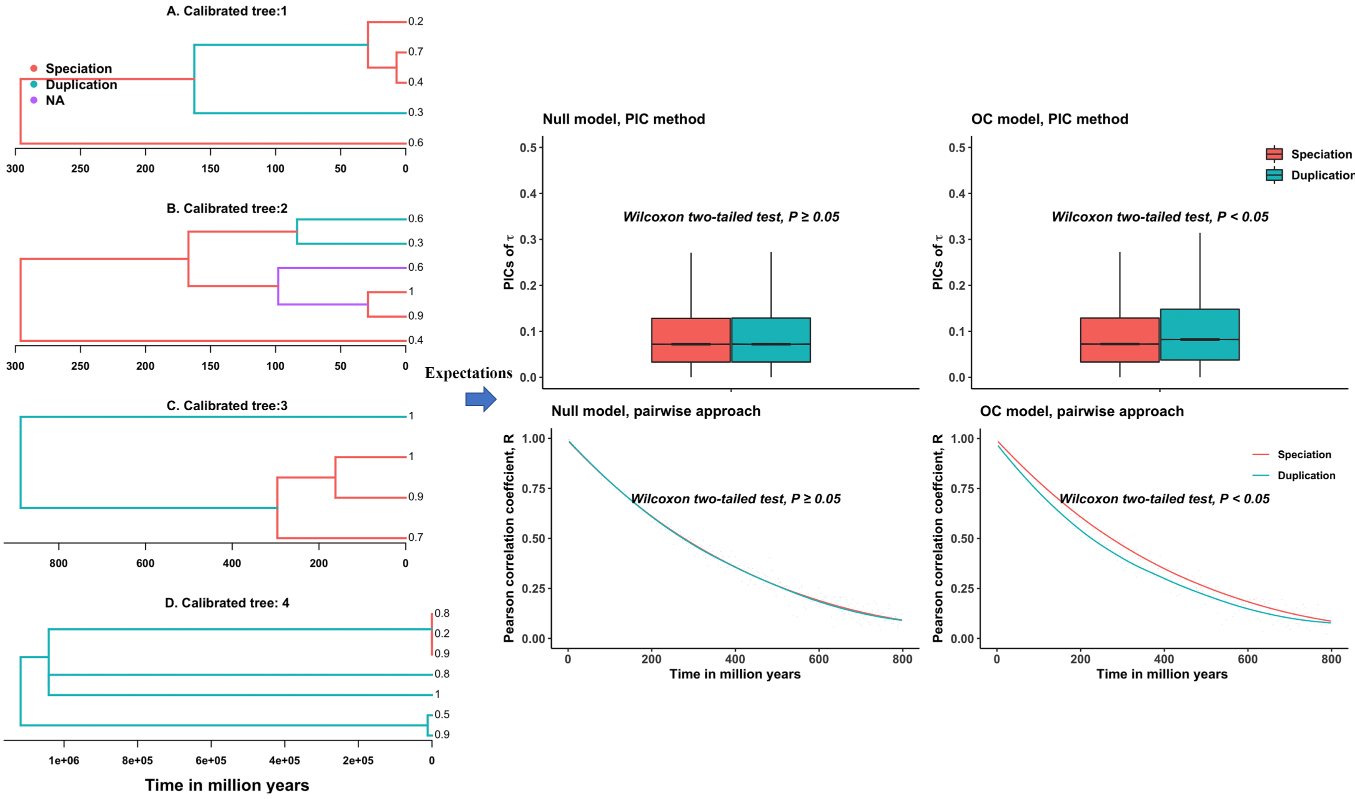

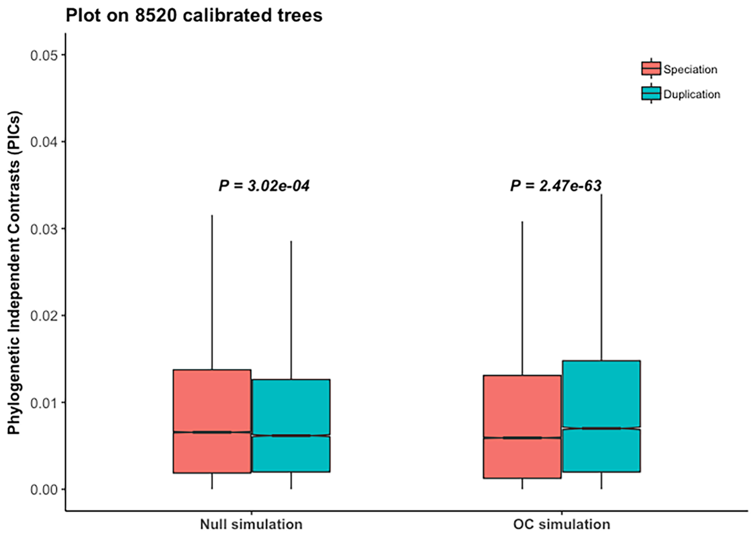

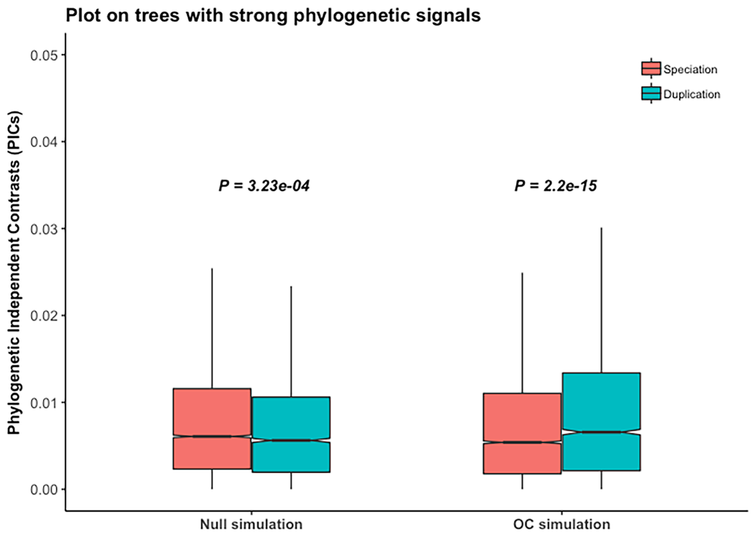

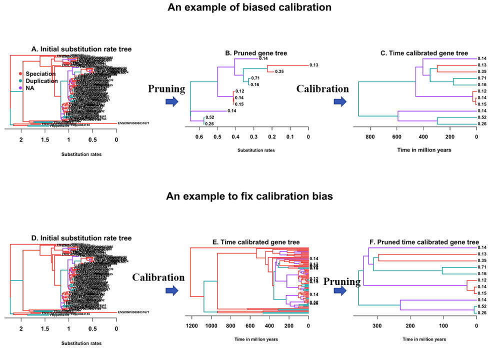

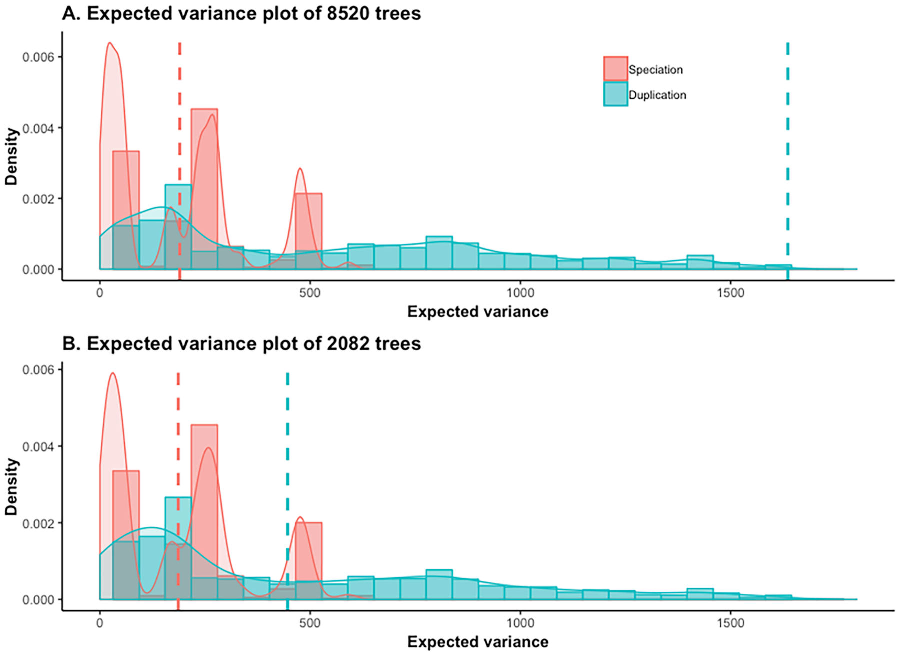

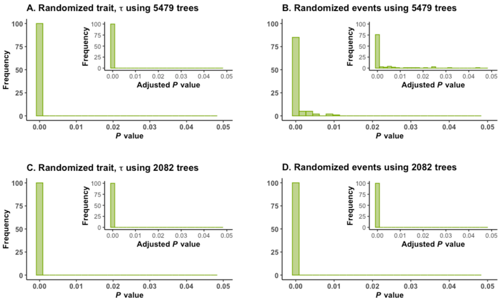

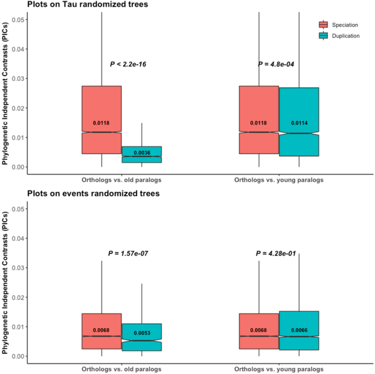

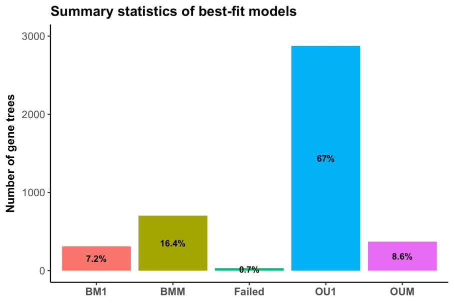

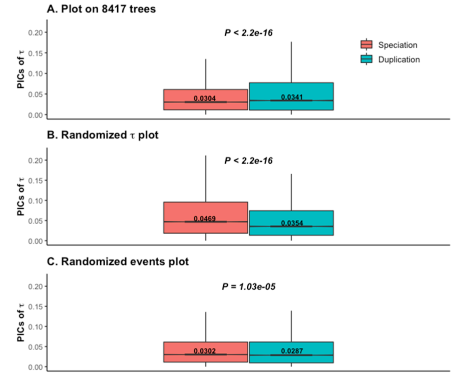

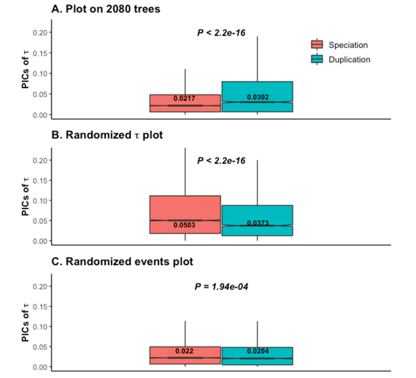
